# The biological activity of bacterial rhamnolipids is linked to their molecular structure

**DOI:** 10.1101/2023.01.23.525263

**Authors:** Bredenbruch Sandra, Müller Conrad, Atemnkeng Henry, Schröder Lukas, Tiso Till, Lars M. Blank, Florian M.W. Grundler, A. Sylvia S. Schleker

## Abstract

Biosurfactants are amphiphilic compounds of microbial origin with a wide range of industrial applications. Rhamnolipids (RLs) offer a broad range of potential applications as biosurfactants in industry and agriculture. Several studies report a high capacity for controlling different plant pests and pathogens and consider RLs as promising candidates for bio-based plant protection agents. RLs are a class of glycolipids consisting of one (mRLs) or two (dRLs) rhamnose moieties linked to a single or branched β-fatty acid. Due to the diversity of RLs, so far little is known about the relation between molecular structure and biological activity.

Engineering the synthesis pathway allowed us to differentiate between the activities of mixtures of pure mRL and of pure dRL congeners and elaborated HPLC techniques further enabled to analyse the activity of single congeners. In a model system with the plant *Arabidopsis thaliana* and the plant-parasitic nematode *Heterodera schachtii* we demonstrate that RLs can significantly reduce infection, whereas their impact on the host plant varies depending on their molecular structure. While mRLs reduced plant growth even at low concentration, dRLs showed no or a beneficial impact on plant development. In addition, the effect of RLs on plant H_2_O_2_ production was measured as an indicator of plant defense activity. Treatment with mRLs or dRLs at a concentration of 50 ppm increased H_2_O_2_ production. Lower concentrations of up to 10 ppm were used to stimulate plants prior to a treatment with water or flagellin (flg22), a bacterial inducer of plant defense responses. At 10 ppm both mRLs and dRLs fostered an increased response to flg22. However, mRLs also led to an increased response to water, the non-inducing negative control, indicating a generally elevated stress level. Neither mRLs nor dRLs induced expression of plant defense marker genes of salicylic acid, jasmonic acid and ethylene pathways within a 1 hour and 48 hours treatment.

Due to the negative effect of mRLs on plants further studies were concentrated on dRLs. Treatment of pre-parasitic infective juveniles of *H. schachtii* revealed that dRLs did not increase mortality even at a very high concentration of 755 ppm. In order to analyse the effect of single dRL congeners nematode infection assays were performed. While dRL congeners with a C10-C8 acyl chain increased nematode infection, dRLs with C10-C12 and C10-C12:1 acyl chains reduced nematode infection even at concentrations below 2 ppm. Plant growth was not reduced by C10-C8 dRLs, but by C10-C12 and C10-C12:1 dRLs at concentrations of 8.3 ppm. H_2_O_2_ production was increased compared to the water control upon treatment with C10-C8 dRLs at a concentration of 200 ppm, while C10-C12 and C10-C12:1 dRLs triggered the same effect already at 50 ppm.

Our experiments show a clear structure-effect relation. In conclusion, functional assessment and analysis of mode of action of RLs in plants require careful consideration of their molecular structure and composition.

## Introduction

Rhamnolipids (RLs) are glycolipidic biosurfactants produced as secondary metabolites by different bacteria of which *Pseudomonas* spp. and *Burkholderia* spp. are studied best. RL synthesis is regulated by quorum sensing in a cell-density dependent manner and synchronises with the stationary phase of bacterial growth [1]. A set of three genes, namely *rhlA, rhlB* and *rhlC*, is responsible for the final synthesis steps from the fatty acid precursor 3-(3-hydroxyalkanoyloxy)alkanoic acids (HAAs) to mono-RLs (mRLs) and eventually to di-RLs (dRLs) [2]. The lipophilic moiety HAA plays a pivotal role in the structural diversity of these molecules via variation in chain length, saturation and the number of fatty acid chains attached to either one or two rhamnose sugars. Until today, there are about 110 congeners described and the congener range and composition depends on the producer strain and on external stimuli such as nutritional parameters [3–5]. Hence, variation in RL synthesis mirrors environmental requirements and challenges and the specific RL composition shapes its nature as a specific adjustment to environmental conditions. It is assumed that the competency of a RL mixture mainly relies on the properties of the most abundant congener [4]. Indeed, the impact of the structure and size of individual congeners is demonstrated for tensioactive traits like the critical micelle concentration (CMC) [4]. The CMC of RLs ranges from 5-200 ppm and is strongly influenced by the ratio of mRLs and dRLs [6]. Information on the influence of the size of the head group on micelle formation is inconsistent though [6,7]. The biological role of RLs is usually associated to their tensioactive properties contributing to nutrient availability, biofilm creation and bacterial motility, but it also involves quorum sensing and antimicrobial activity [8,9]. In particular, the numerous reports of the antimicrobial activity of RLs indicate an outstanding potential of these bacterial molecules e.g. as biologicals in environmentally friendly crop protection [3]. The question regarding their mode of action still needs clarification and, by all appearances, the answer combines multiple aspects. Antioomycete activity of RLs is often associated with direct lysis of zoospores due to the absence of a cell wall while antibacterial activity is less radical and proposed to rely on cell permeabilization and changes in cell hydrophobicity [10,11]. Several studies describe plant immune stimulation by RLs [12–14], but the individual steps starting with the nature of RL perception to early signaling processes are still a matter of speculation. One crucial obstacle in understanding the mode of action is the lack of comparability between studies due to insufficient definition or differences in RL compositions. So far, the effect of individual RL types and congeners has been largely neglected although obviously affecting the biophysical character and likely biological activity. Therefore, we analysed the efficacy of different RLs to impair the host-parasite interaction in a model system with *Arabidopsis thaliana* and the plant-parasitic nematode (PPN) *Heterodera schachtii*. As a cyst nematode, this species is among the economically most important PPNs, and in addition, an eminent model to investigate parasitic interactions of PPNs with plants [15,16]. Successful RL mixtures were further analysed with focus on their influence on plant immunity including the expression of defense markers and their role as direct and indirect elicitors of oxidative signaling. Eventually, the effect of the most efficient RL type was evaluated on congener level.

## Results

### Effect of RLs on nematode infection and plant growth

The effect of RLs on *H. schachtii* infection was validated with mixtures of HAAs, mRLs or dRLs supplemented to the sterile plant growth medium. RLs clearly affect nematode infection at a concentration of only 8.3 ppm (parts per million, equals mg/l), whereas the same concentration of HAAs is ineffective. Thus, it requires the complete biosurfactant molecule comprising dTDP-L-rhamnose and HAAs to significantly reduce the number of adult nematodes per plant (Fig. 1a). However, no clear adverse effects in female size development and nematode reproduction could be observed (Fig. 1b and 1c). Furthermore, the number of dTDP-L-rhamnose of the RL molecule considerably impacts host plant development. A mRL concentration of only 8.3 ppm lowers root and shoot growth to a minimum, while the same concentration of HAAs reduces only root growth by about 15%. In contrast, dRLs increase root growth by about 15% and simultaneously reduce the infection by about 75% (Fig. 1d, e). This experiment shows that mRLs, dRLs, and HAAs clearly have different effects on plant and nematode development. As HAAs do not impact nematode infection, we then focused on analysing the effect of mRLs and dRLs on plant H_2_O_2_ production.

**Figure 1.**
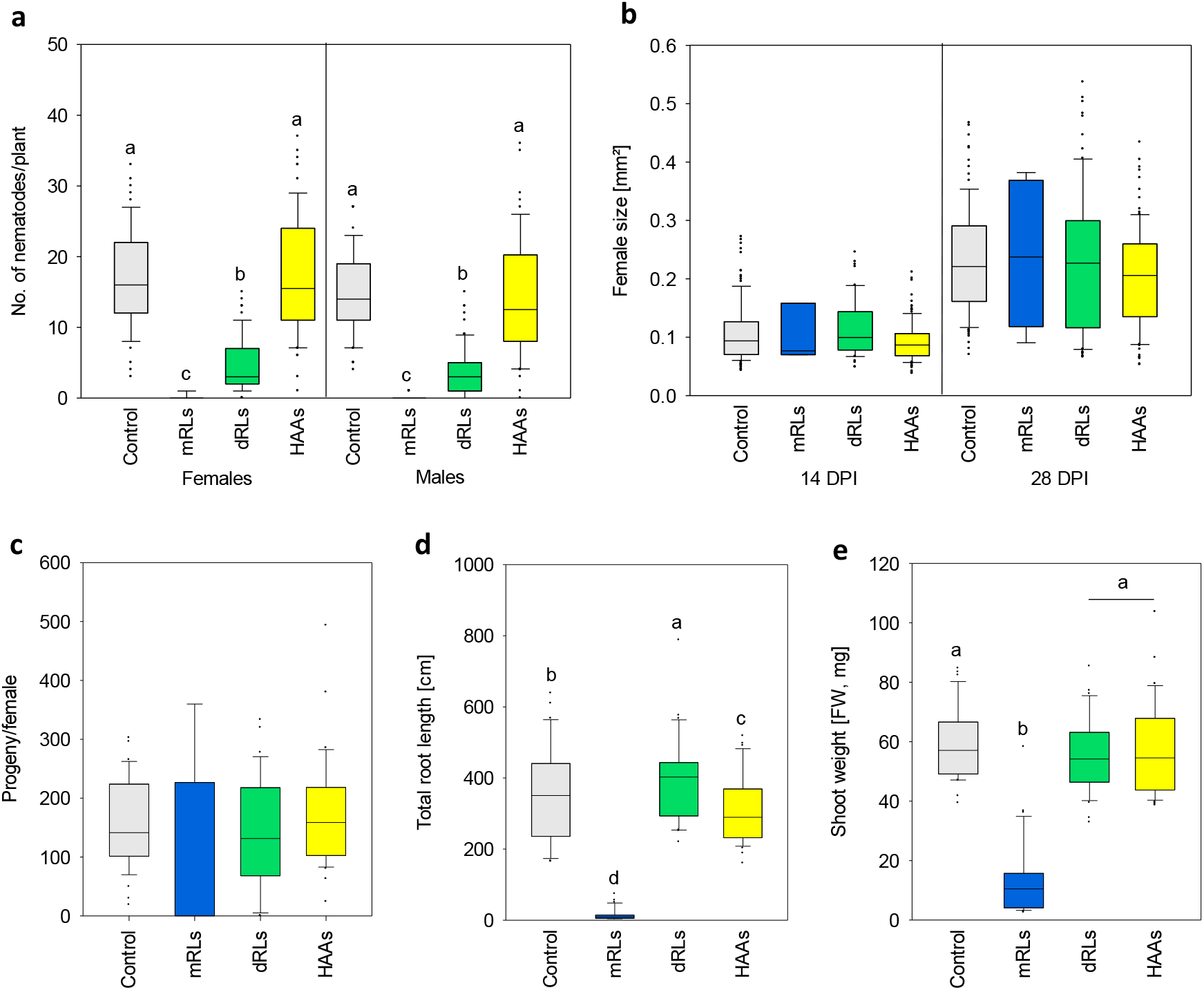
Impact of different RLs and HAAs on nematode parasitism and host plant A. thaliana inoculated with H. schachtii under in vitro conditions. The compounds were supplemented to the plant growth medium in a concentration of 8.3 ppm and their impact on a) the number of adult nematodes at 14 days post inoculation (dpi), b) the size of female nematodes at 14 and 28 dpi, and c) progeny per female at 35 dpi, d) total root length (cm) and e) shoot fresh weight (mg) at 14 DPI was analysed. Median with 25%- and 75%-quartile of three independent biological replicates with total n=70 plants is shown for the nematode data and total n=34 plants for the plant data. Significance is based on ANOVA on ranks and post-hoc Student–Newman–Keuls (SNK), different letters indicate significant differences between treatments (p<0.05).

### Effect of mRLs and dRLs on H_2_O_2_ synthesis of *A. thaliana* induced by flg22

Production of extracellular H_2_O_2_ is one of the earliest plant responses upon pathogen infection or stimulation by pathogen-associated molecular patterns (PAMPs). Therefore, we tested whether RLs are able to modify H_2_O_2_ production in leaves. Concentration levels were elevated to 50 and 200 ppm to ensure that a potential plant response was not below detection level. Increasing H_2_O_2_ production could be observed upon exposure to both mRLs and dRLs compared to water. Interestingly, for both RL types total H_2_O_2_ production was generally higher with 50 ppm RLs and only for mRLs, it was also significantly increased by treatment with 200 ppm (Supplementary Fig. 2). Additionally, we analysed the expression of several defense-related genes upon root exposure for 1 hour and after 48 hours to 8.3 ppm mRLs and dRLs. The later time point displays the state of plant defense J2s encountered in the infection assays after inoculation. Neither a treatment with mRLs nor with dRLs results in remarkable variation in the expression of selected marker genes compared to the control (Supplementary Fig. 3).

Since RLs obviously do not act as primary PAMPs, we subsequently tested, whether they are able to modulate the plant’s immune response in advance to PAMP exposure. Therefore, H_2_O_2_ synthesis in leaves was measured in response to the well-described PAMP flagellin 22 (flg22) after a pre-treatment with different concentrations of mRLs or dRLs (Fig. 2). Leaves were harvested and incubated overnight in different concentrations of mRLs, dRLs or water as control. The following day, RLs were washed off and H_2_O_2_ levels in response to ddH_2_O or flg22 were measured.

**Figure 2.**
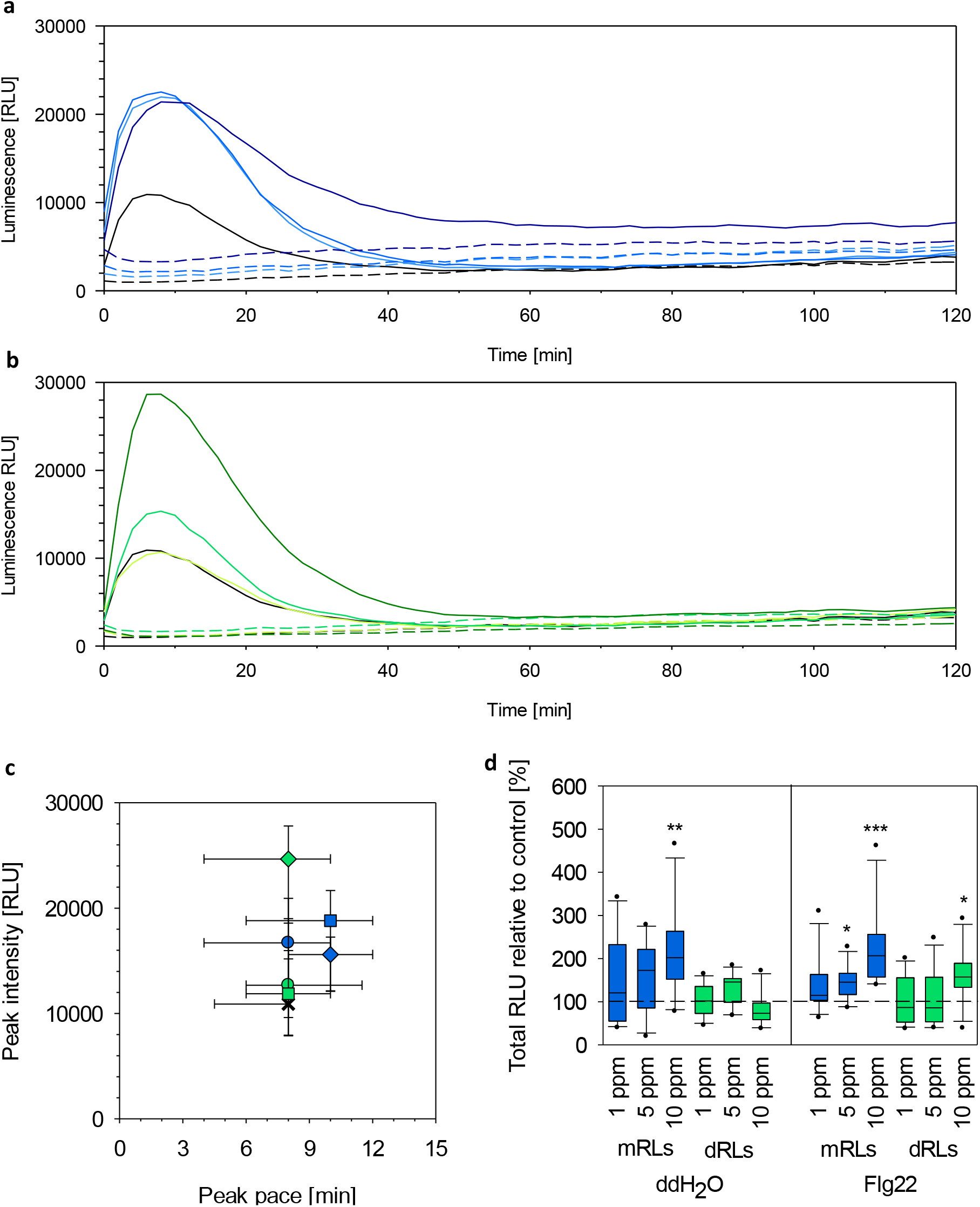
Impact of different RLs on plant H_2_O_2_ production induced by flg22. Arabidopsis thaliana leaves were incubated in mRLs (blue) and dRLs (green) solutions at different concentrations and subsequent synthesis of H_2_O_2_ induced by the bacterial elicitor flg22 was monitored over a period of two hours. a) H_2_O_2_ production over time after mRL incubation, dashed lines: response to ddH_2_O, solid lines: response to flg22, black: mock control, from light to dark blue illustrates an increase in concentration (1, 5, 10 ppm) of mRLs used for stimulation, mean values; b) H_2_O_2_ production over time after dRL incubation, dashed lines: response to ddH_2_O, solid lines: response to flg22, black: mock control, from light to dark green illustrates an increase in concentration (1, 5, 10 ppm) of mRLs used for stimulation, mean values; c) Time and intensity of H_2_O_2_ peak induced by flg22, cross: mock control, circle: 1 ppm, square 5 ppm, diamond: 10 ppm, median with 25%- and 75%-quartile; d) Total H_2_O_2_ produced within the initial 2 hours of flg22 induction, median with 25%- and 75%-quartile, significance based on 2-tailed t-test compared to water control (dashed line) with * p<0.05, ** p<0.01, *** p<0.001. a)-d) data of three independent biological replicates with total 11≤n≤12.

Both RL types led to an increased leaf H_2_O_2_ production upon elicitation with flg22 compared to the mock control. Here, mRLs induce significant increase already at 5 ppm whereas a concentration of 10 ppm is required to have a similar effect after pre-treatment with dRLs (Fig. 2d). Although the overall H_2_O_2_ production after stimulation with 10 ppm mRLs is doubled compared to the mock control, the highest peak intensity of the flg22-induced curve is observed after stimulation with 10 ppm dRLs (Fig. 2c). A major difference though is the H_2_O_2_ synthesis following this peak. After dRL stimulation, it quickly declines to the level of the water control whereas after mRL stimulation, the decline is less intense and remains a multiple of the water control for the whole time monitored (Fig. 2a,b). Another major difference after pre-treatment with mRLs and dRLs is the basal H_2_O_2_ production in response to negative control. Water is non-stimulatory, still the H_2_O_2_ production is double as high after stimulation with 10 ppm mRLs and a similar trend also for 5 ppm mRLs (Fig. 2c). In contrast, there is no change in H_2_O_2_ production after stimulation with any concentration of dRLs. Both RL types modify the H_2_O_2_ profile towards flg22 for a stronger reaction, but in contrast to dRLs a pre-treatment with mRLs also elevates the basal H_2_O_2_ level. Although mRL treatment reduces nematode infection, plants are easily stressed and show strong growth decline (Fig. 1d-e, 2d). Therefore, it is possible that the effect observed on plant susceptibility to *H. schachtii* results from deteriorated host plant qualities due to a decrease in root mass. In contrast, dRLs impair an infection without phenotypical disadvantages for the host plant. Thus, further experiments were conducted with dRLs only.

### Nematicidal activity of dRLs

A mortality assay with the nematode in its pre-parasitic stage was performed in order to understand whether a potential lethal impact of dRLs could be responsible for the observed infection decline. Exposure of J2s to dRL concentrations in the range used in the infection assay was noneffective (data not shown). Even concentrations inordinately high had no nematicidal effect in a range 45 times as high as used in the infection assays (Fig. 3). Hence, it seems improbable that the decrease in the number of infected nematodes results from direct nematicidal activity of dRLs.

**Figure 3.**
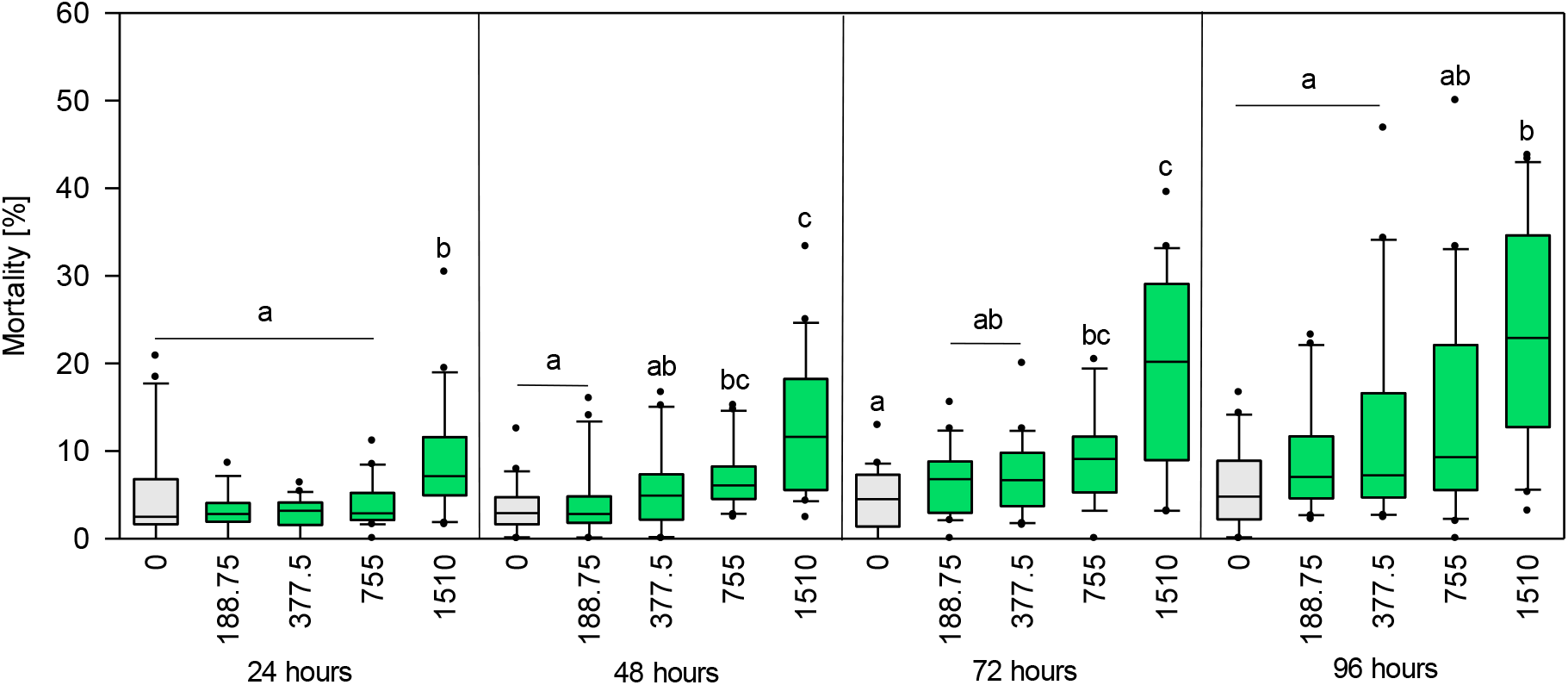
Effect of dRLs on mortality of H. schachtii infective stage juveniles (J2). Nematodes were kept in solutions of dRLs at different concentrations for a period of four days. Median with 25%- and 75%-quartile of three independent biological replicates with in total n=20 wells (about 50 J2/well) per treatment and time point. Significance based on ANOVA on ranks and post-hoc Dunn’s Method, different letters indicate significant differences between treatments (p<0.05).

### Effect of single dRLs congeners on nematode infection and plant

As a final step for understanding the structural importance of RLs, the impact of single dRL congeners on nematode infection and plant H_2_O_2_ production was analysed. Each dRL congener was analysed in two different concentrations. The first concentration was chosen to compare the efficacy of all congeners at the same concentration (Fig. 5). The second concentration was chosen following the hypothesis of the most dominant congener directing the overall character of the mixture (Fig. 6). Here, each congener was supplemented based on the proportion given in the dRL mixture used in the previous experiments at 8.3 ppm: 5% C10-C8 dRL, 61% C10-C10 dRL, 19% C10-C12 dRL, 14% C10-C12:1 dRL.

**Figure 4.**
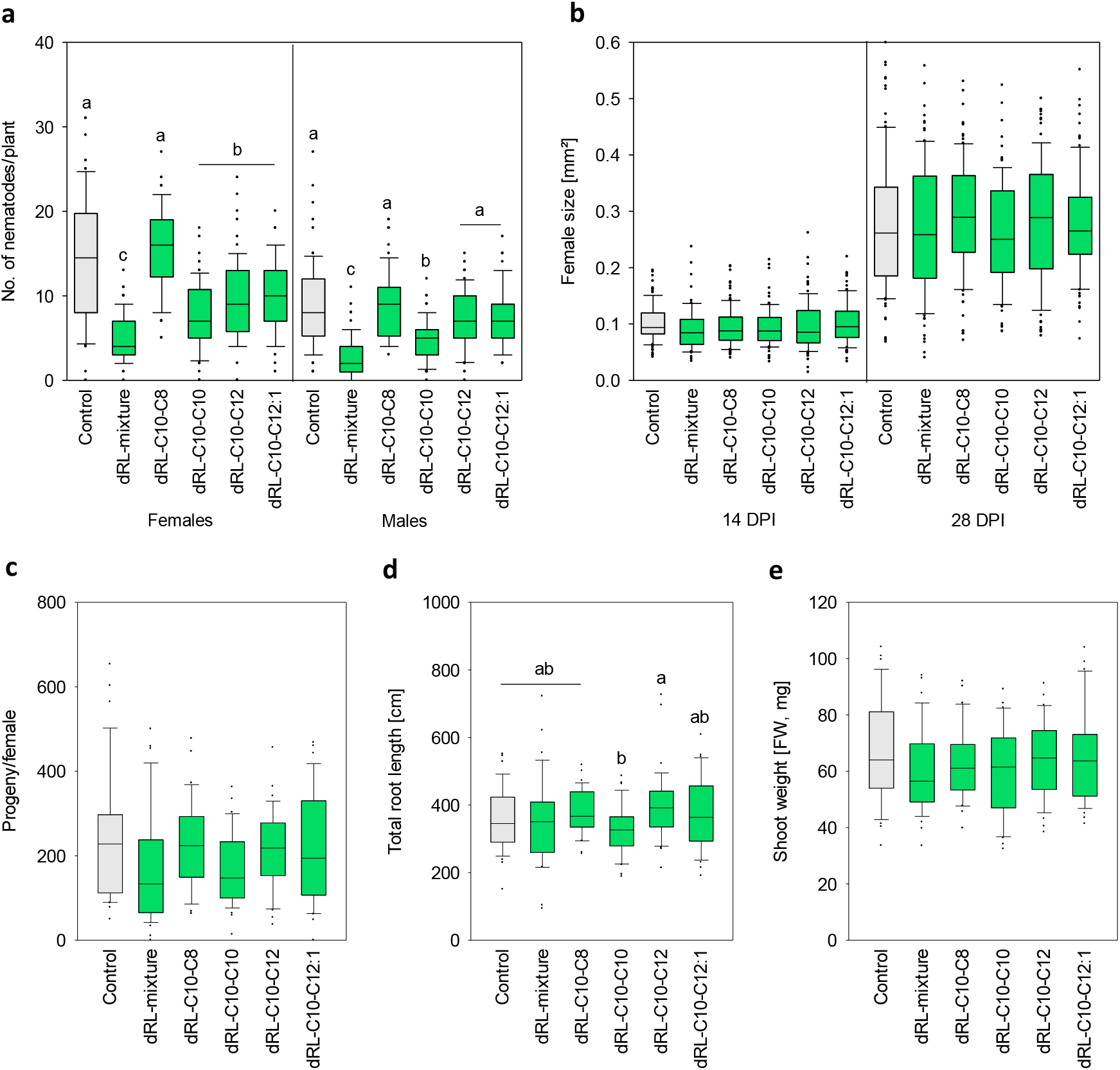
Impact of di-rhamnolipid (dRL) congeners in their relative percentage in a natural dRL mixture at 8.3 ppm as plant growth medium supplement (0.4 ppm Rha-Rha-C10-C8, 5.1 ppm Rha-Rha-C10-C10, 1.6 ppm Rha-Rha-C10-C12, 1,2 ppm Rha-Rha-C10-C12:1) on the plant and infection phenotype of Arabidopsis thaliana infected with Heterodera schachtii. Displayed are the number of adult nematodes at 14 days post inoculation (DPI), the size of females at 14 and 28 DPI, and progeny per female at 35 DPI for the infection phenotype and the total root length (cm) and shoot fresh weight (mg) at 14 DPI for the plant phenotype. Median with 25%- and 75%-quartile of three independent biological replicates with total n≥64 plants for the infection phenotype and total n≥32 plants for the plant phenotype. Significance based on ANOVA on ranks and post hoc Dunn’s Method, different letters indicate significant differences between treatments (p<0.05).

**Figure 5.**
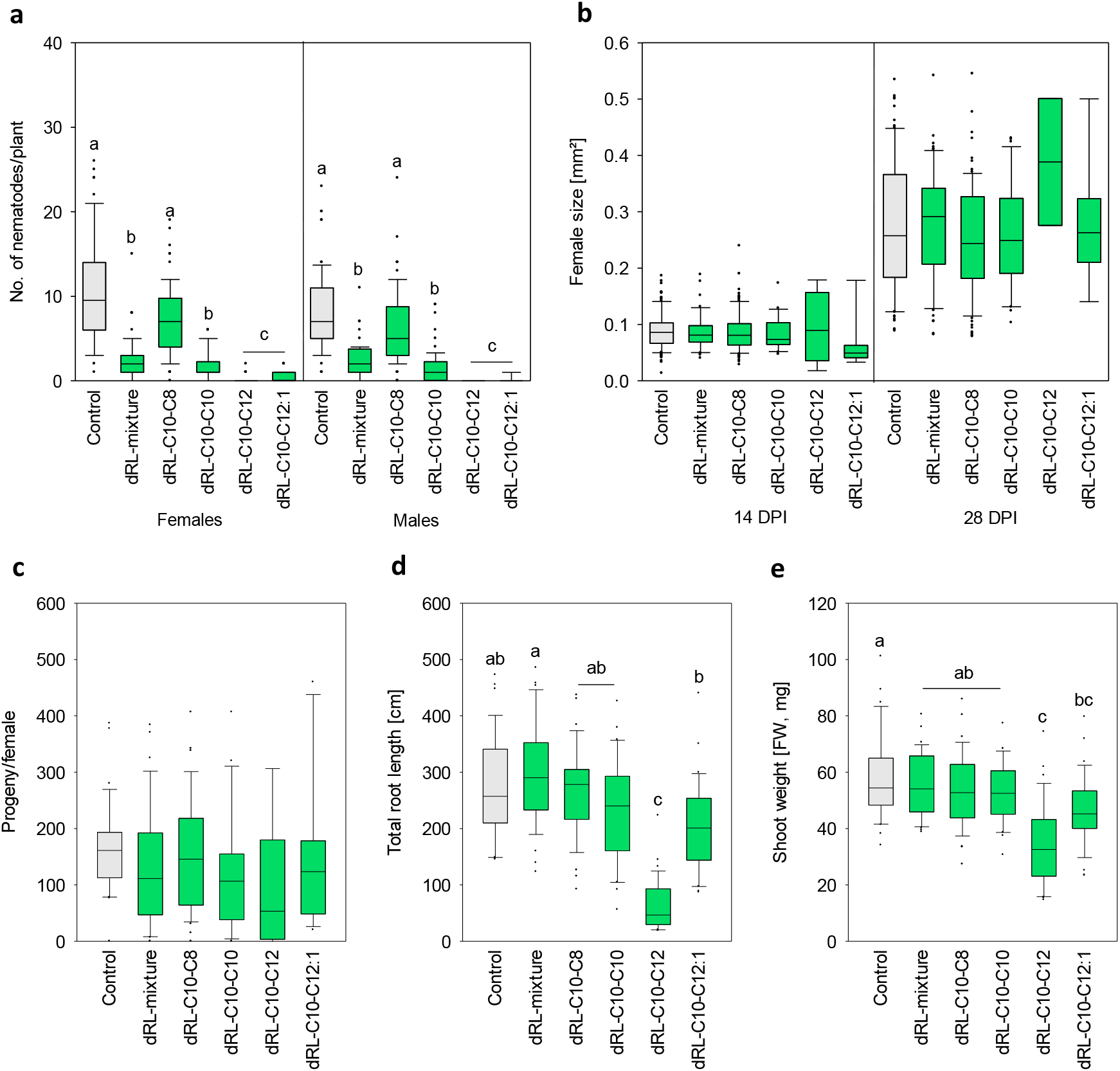
Impact of di-rhamnolipid (dRL) congeners as 8.3 ppm plant growth medium supplement on the infection and plant phenotype of Arabidopsis thaliana inoculated with Heterodera schachtii. Displayed are the number of adult nematodes at 14 days post inoculation (DPI), the size of females at 14 and 28 DPI, and progeny per female at 35 DPI for the infection phenotype, and the total root length (cm) and shoot fresh weight (mg) at 14 DPI for the plant phenotype. Median with 25%- and 75%-quartile of three independent biological replicates with total n≥66 plants for the infection phenotype and total n≥32 plants for the plant phenotype. Significance based on ANOVA on ranks and post hoc Dunn’s Method, different letters indicate significant differences between treatments (p<0.05).

**Figure 6.**
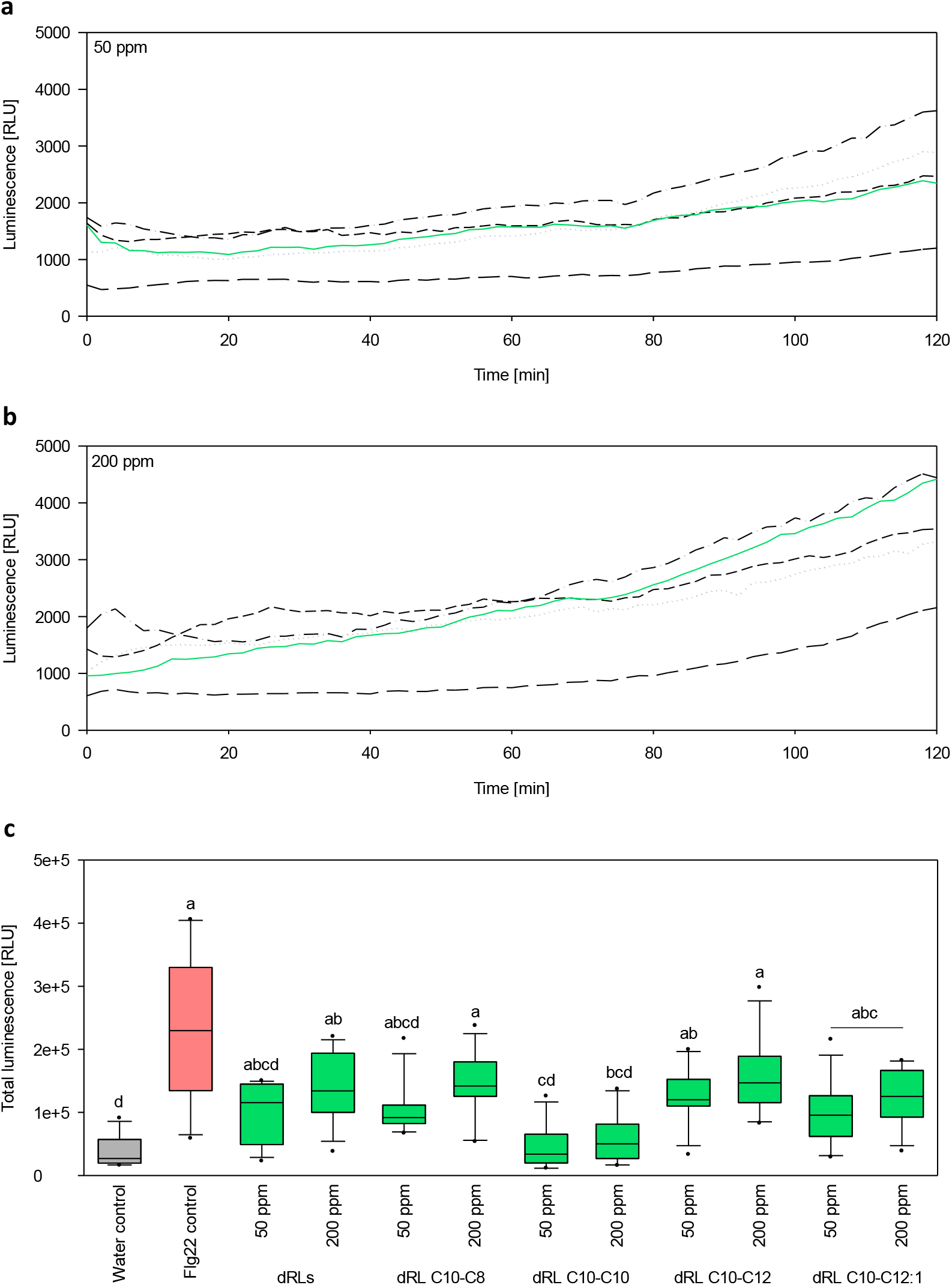
H_2_O_2_ response of Arabidopsis thaliana leaf tissue to different concentrations of single di-rhamnolipids (dRLs) congeners and the original dRL mixture. (a) H_2_O_2_ over a period of two hours. Median with 25%- and 75%-quartile of three independent biological replicates with total 10≤n≤12. (green: dRLs, small dash: dRL C10-C8, long dash: dRL C10-C10, dash-dot: dRL C10-C12, dotted: dRL C10-C12:1) (b) Total H_2_O_2_ after two hours. Mean of total 10≤n≤12 of three independent biological replicates. Significance based on ANOVA on ranks and post hoc Dunn’s Method, different letters indicate significant differences between treatments (p<0.05).

None of the single dRL congeners can entirely recreate the effect induced by the dRL mixture, but aside from C10-C8 dRLs all of them are able to interfere with nematode establishment at the root. The efficacy depends not only on the respective concentration, but correlates with the 3-OH acyl chain length. RLs with longer chains appear to be more efficient at lower concentrations. Very low concentrations of below 2 ppm of C10-C12 and C10-C12:1 dRLs reduce the infection as effective as the predominant congeners C10-C10 dRL with more than 5 ppm (Fig. 5a). Notably, root and shoot growth is significantly impaired during root exposure to 8.3 ppm of these longer dRL congeners (Fig. 6d and 6e). This is further confirmed by the H_2_O_2_ profile of single dRL congeners (Fig. 7). Already 50 ppm of C10-C12 and C10-C12:1 dRLs cause a significant increase in H_2_O_2_ production compared to the water control. A comparison between the dRL mixture and its single congeners reveals no difference in the total H_2_O_2_ synthesis, but in the H_2_O_2_ pattern. Exclusively upon exposure to C10-C12 dRLs there is a peak at 200 ppm within less than ten minutes while the mixture and all other single congeners induce a steady, but unspecific increase in H_2_O_2_ over time (Fig. 8c).

Hence, the individual dRL congeners differ in their impact on nematode infection and the plant in comparison to each other as well as to the complete mixture with a specific responsiveness towards longer dRL congeners.

## Discussion

RL biosynthesis involves a mixture of different RL congeners generating a high structural diversity. Although the general RL profile is usually similar among one bacterial species the exact RL composition varies widely. The ratio of mRLs and dRLs and the relative abundance of minor congeners is influenced by the producer strain and the media composition [6]. Changes in the ratio of mRLs and dRLs and the acyl chain length are known to affect the tensioactive properties of the RL mixture, such as the CMC [4,17]. However, the structure-dependent activity of RLs is still uncertain when it comes to their ability to counteract pathogens.

Our data show that dRLs are potent agents to reduce PPN development in plant without direct lethal impact on the nematodes. On the plant side, we observed immune-related resposes that may contribute to plant defense against the PPN. RLs triggered an enhanced oxidative level and boosted H_2_O_2_ synthesis upon exposure to the pathogenic stimulus flg22. Interestingly, plant exposure to RLs was not accompanied by an altered defense gene expression.

Moreover, the effect on the plant side, and thus the potential to control nematodes, highly depends on the type of RL with additional specificity on congener level. These observations are essential for a comprehensive understanding and eventual application of RLs for crop protection purposes.

### Effect of dRLs on nematode parasitism

We found that dRLs have no direct nematicidal activity. However, even at low concentration of dRLs in the growth substrate, parasitism of nematodes is significantly inhibited. Change in sex determination, retarded development of females and reduced fecundity are typical effects occurring in resistant plants. However, these effects could not be observed. Therefore, we assume that dRLs interfere with early events of nematode parasitism.

One key step in early parasitism events is nematode migration towards the host plant rhizosphere. Root exudates play a crucial role to attract nematodes and guide their way to the host plant [18]. Zhao et al. found that J2s of the root-knot nematode *Meloidogyne incognita* did no longer accumulate in the area of the root tip of pea roots when border cells and thus their root exudates were washed off with water before nematode inoculation [19]. Considering the tensioactive character of surfactants, RLs are even more potent agents to detach root compounds to alter the characteristic root exudate composition required for nematode orientation. Interference with nematode orientation could also be linked to a peculiarity of RLs closely resembling ascarosides. These glycolipidic nematode pheromones comprise the dideoxysugar ascarylose linked to a fatty acid-like side chain and play a crucial role in basal nematode activities [20]. Recently, it was observed that *A. thaliana* is able to absorb and process ascarosides turning them into a repellent for PPNs [21]. It is a matter of further studies to find out, whether this is also the case for RLs. In 1989, Terzaghi successfully used the synthetic surfactant Tween as conjugate to transfer exogenous fatty acids into different plant species [22]. In similar experiments with *A. thaliana*, Tween detergents enabled the uptake of polyhydroxyalkanoates, but the efficiency was dependent on the type of Tween indicating that there are attributes in addition to the amphiphilic nature that determine the success of this process [23]. If bacterial RLs can pass the plant membrane like their synthetic counterpart Tween or their nematode-derived structural equivalent ascaroside a modification of the RL molecules as described for ascarosides may similarly affect nematode orientation outside or inside plants and thus impair nematode parasitism in particular at an early stage.

Another key step during early parasitism is host invasion and the induction of the feeding site. At this stage, host plant defense is highly active. However, neither ethylene-, nor jasmonic acid- or salicylic acid-related defense markers showed altered expression levels in response to 8.3 ppm RLs. Similar observations with different RL mixtures, types and concentrations can be found in literature. Regulation of the tomato defense markers *PR1* and *PRP69* was not affected 3 days after stimulation with 300 μM synthetic mRLs [14]. In grapevine cells, 10 ppm of a mixture of 40% C10-C10 mRLs and 60% C10-C10 dRLs induced upregulation of the *chit4c* defense marker after 9 hours [12]. In contrast, 200 ppm of the same composition had no effect on the expression of *PR1, PR4* and *PDF1.2* in *Arabidopsis* leaves 48 hours post treatment [13]. A significant upregulation was only found for *PR1* sprayed with 1000 ppm of the RL mixture. Apparently, plant defense gene activation induced by RLs is possible, but not necessarily the prevalent response. The plant species and tissue type play a role just like the RL concentration, however, it is debatable how to interpret plant defense activation induced by biological compounds at concentrations far beyond their natural purpose.

Data on the impact of RLs on plant H_2_O_2_ synthesis was more conclusive. Incubation in 10 ppm RLs led to an elevated flg22-induced H_2_O_2_ amplitude and general H_2_O_2_ level in comparison to the mock control. Perception of the bacterial elicitor flg22 entails an oxidative burst that stimulates plant immune activity [24,25]. An increase in H_2_O_2_ synthesis as we observed in response to flg22 after RL treatment may reinforce signalling and eventually plant defense. Therefore, we assume that rather than directly activating plant defense gene expression, RLs prime the immune system for reinforced plant defense upon a second trigger.

Our data indicate that RLs interfere with nematode infection in the early stages of parasitism. Based on our observations together with literature, we suggest two possibilities how RLs affect nematode infection: interference with nematode orientation and/or plant defense priming.

### RLs as priming agents against PPNs

According to Martinez-Medina et al., defense priming is a two-step mechanism initiated by a priming stimulus followed by a stressor as the trigger stimulus [26]. Plant fitness benefits from the primed stage in two ways: upon pathogen attack plant defense activation is more robust while plant maintenance costs are lower in comparison to unprimed defense activation. Correspondingly, in our infection assays we observed that during dRL treatment nematode infection is reduced while root growth is better under conditions of infection pressure. Interestingly, mRL treatment strongly impairs plant growth even without pathogen pressure (data not shown).

Based on literature, defense priming activity induced by RL depends on the trigger stimulus. Stimulation of grapevine cells with 5 ppm of a mixture of 40% C10-C10 mRLs and 60% C10-C10 dRLs led to an increased oxidative burst induced by chitosan, but not by *Botrytis cinerea* [12]. However, stimulation with 200 ppm of the same RL mixture potentiated the expression of *PR1* in *Arabidopsis* after inoculation with *B. cinerea* 4 days later, while neither *PR1* or *PDF1.2* were further upregulated when inoculated with *Pseudomonas syringae* pv. *tomato* DC3000 or *Hyaloperonospora arabidopsidis* [13]. Potentiated expression of *PR1* and of *PRP69* in response to *B. cinerea* was also confirmed in tomato after stimulation with 300 μM synthetic mRLs [14].

Defense priming is further associated with several adjustments that allow plants to react faster and/or stronger to stressors. One adjustment is the accumulation of pattern recognition receptors (PRRs) such as the transmembrane receptor kinase FLAGELLIN SENSITIVE2 (FLS2) which regulates the perception of flg22 [25,26]. Our data show a boost in H_2_O_2_ synthesis after RL stimulation, but it is unclear if this effect is related to FLS2 accumulation Future work should further focus on the expression level of PRRs after RL treatment, including PRRs for PAMPs like FLS2 as wells as for nematode-associated molecular patterns (NAMPSs). The field of NAMPs is still relatively new, but it was observed that plant reaction to NAMPs, PAMPs and microbe-associated molecular patterns (MAMPs) is similar suggesting that plants have evolved receptors specific for nematode perception [27,28]. Here, the *Arabidopsis* leucine-rich repeat receptor-like kinase (NILR1), the first receptor described to be involved in plant immunity to PPNs, would be a suitable candidate [28].

Another adjustment of the plant memory during plant defense priming is histone modification and thus the regulation of transcription factors [26,29]. However, we could not confirm defense priming activity based on transcription factor expression using MYB72. The root-specific transcription factor MYB72 is described to be essential for defense priming activation induced by root colonization of *Arabidopsis* with nonpathogenic *Pseudomonas fluorescens* WCS417r, another *Pseudomonas* species described to produce RLs and thus, a valid candidate in our expression analysis [11,30]. More generally, *AtMYB* genes are involved in secondary metabolism which often requires specific conditions to expedite a phenotype [31]. The activity of *MYB72* is furthermore associated with iron-deficiency and Palmer et al. suggested that the onset of *MYB72*-related priming may rather be linked to a competition for iron [32]. Still, there are further suitable transcription factors, e.g. of the WRKY family, linked to gene priming which should be considered in future experiments to further define priming activity of RLs [33].

Defense priming in plants is confirmed for diverse stimulants, plenty of them related to bacteria. Bacterial dRLs meet some of the criteria described by Martinez-Medina et al. to confirm including low fitness costs and more robust plant defense [26]. However, the memory effect of the priming stimulus is only indicatively addressed in our experiments. Aspects like level of PRRs, in particular for the perception of PPNs, and the expression of certain transcription factors require more research. RL perception in plants is still not comprehensively understood, but the predominant theory refers to the insertion of RLs into the phospholipid bilayer of membranes influenced by the structure and size of RLs [34–37] omitting the possibility of transmembrane transfer. Together with or in addition to membrane intercalation, RLs may disrupt the epidermal membrane causing detachment of plant-derived molecules. The plant surface is equipped with a variety of PRRs specified to perceive self-molecules during wounding called damage-associated molecular pattern (DAMP) receptors [38]. Although the underlying RL perception mechanism in plants is still unraveled, we demonstrate that RLs are triggers of plant defense whereby quality and intensity of plant responses depend on the concentration and structure of RLs.

### *A. thaliana* differs in its sensitivity towards different RL congeners

Many studies in the field of pathogen control using RLs refer to mixtures comprising both mRLs and dRLs with *P. aeruginosa* as the main producer species. There are relatively few studies that involve the structural diversity of RLs, often with inconsistent results. Sha et al. found that fungicidal activity was more powerful for dRLs compared to mRLs (produced by *P. aeruginosa* ZJU211) against the two *Oomycetes Phytophthora infestans* and *P. capsici*, three *Ascomycota B. cinerea, Fusarium graminearum* and *F. oxysporum* and two *Zygomycetes Mucor circinelloides* and one *Mucor* spp. [39]. In other publications, it is mRLs (produced by *P. aeruginosa*) that were more effective against *Phytophthora sojae, Alternaria alternata* G2, *Cladosporium* sp. B, *Actinomucor* sp. Y and *Penicillium oxalicum* S11, and even against certain bacteria such as *Escherichia coli* DH5α, *Bacillus wiedmannii* H238, *B. Safensis* B36# and *Pantoea agglomerans* B10 [40,41]. A comparison of mRLs and dRLs from different producers showed that defense gene expression induction in grapevine cells is highly individual in response to the same concentration of mRLs or dRLs from *P. aeruginosa* (predominant C10-C10 acyl chain) or dRLs from *B. plantarii* (predominant C14-C14 acyl chain) [12]. To our knowledge, there are no comparative studies on the effect of RL congeners on plants.

We observed that growth of *A. thaliana* was inhibited in presence of mRLs whereas dRLs caused no unfavorable effect in plant phenotype. This disparity is also reflected in early plant signaling processes revealing a much stronger oxidative response to 50 ppm mRLs than to dRLs. Furthermore, stimulation with only 10 ppm mRLs increased the basal H_2_O_2_ level in response to the non-stimulative water control. Reactive oxygen species (ROS) like H_2_O_2_ are involved in several fields of signal transduction including the hormonal crosstalk that regulates plant growth [42]. It is suggested that there is a trade-off between stress-related ROS synthesis and vegetative growth describing a shift of resources to either provide plant growth and reproduction or to invest into stress resistance [43]. The poor plant phenotype we observed in the presence of mRLs may result from an imbalance of the natural ROS homeostasis caused by the upregulation of the basal oxidative state.

Pure dRLs demonstrate good potential to suppress parasitism of the PPN *H. schachtii*. Interestingly, we see clear differences in plant response depending on the individual dRL congener applied. Nematode infection was no longer reduced in the presence of pure C10-C8 dRLs while in the presence of dRLs with longer acyl chains infection was still reduced, but the efficiency strongly differed among dRL congeners. The number of female nematodes was as effectively reduced in the presence of dRLs with a C10-C12 or C10-C12:1 acyl chain at a concentration below 2 ppm as in the presence of C10-C10 dRLs at a concentration above 5 ppm. However, a concentration of 8.3 ppm C10-C12 dRLs caused significant plant growth inhibition for root and shoot tissue while the addition of a single double bond in C10-C12:1 dRLs could avert growth inhibition for shoot tissue. With 61%, C10-C10 dRL was the predominant congener in the utilized mixture. In contrast to the assumption that the most abundant congener determines the overall performance of the mixture, we demonstrate that this congener alone is weaker compared to the dRL mixture in terms of the impact on nematode infection and plant phenotype. Although H_2_O_2_ induction by C10-C10 dRLs was similar to the dRL mixture, it differed significantly from C10-C8 and C10-C12 dRLs. A similar observation was made with tomato leaves and ether-synthetic mRLs with an acyl chain length varying from C4 to C18 inducing either no ROS (C4, C6, C14, C16 and C18), moderate (C8) or strong and long-lasting ROS synthesis (C10 and C12) [14]. Furthermore, there is a positive correlation between synthetic RLs with a C12 acyl chain and the biological activity against the fungus *B. cinerea* on tomato and *Zymoseptoria tritici* on wheat [14,44]. Comparing the diverging effects of single congeners we observed on nematode infection, plant development and early plant signalling, we hypothesize that the effect observed for the dRL mixture is the result from a synergism of individual congeners rather than from one dominating congener. Based on our data, a costly separation of RL mixtures down to congener level is no premise for effective plant protection. We assume that a mixture even including mRLs can still be effective as long as the composition of RL congeners is balanced. Therefore, it is recommended to evaluate RL-induced effects on plants and pathogens explicitly based on the respective congener composition. The effectivity of RLs most likely changes due to the absence, presence or even minor differences in the concentration of RL congeners depending on the producing strain and culture conditions. As a consequence, for a definite assessment of a certain capability such as antimicrobial activity or plant immune stimulation the impact of various RL mixtures should be compared. Further exploration of the natural resources of RL producers together with genetic engineering will be a promising approach to further optimize RL compositions towards individual target applications.

In conclusion, RLs are produced with great structural diversity and their effect on plant growth and plant infection with *H. schachtii* differs widely depending on the molecular structure including variation in the hydrophilic as well as in the hydrophobic moiety. Nematode infection could be reduced in the presence of both mRLs and dRLs, but mRL treatment caused plant growth impairments. The effect of dRLs varied on congener level and revealed high efficiency of dRLs with C10-C12 and C10-C12:1 acyl chains to reduce nematode infection. The impact of dRLs on the infection cannot be linked to nematicidal activity, but to the stimulation of plant defense priming. Here, transcriptome sequencing may enable to elucidate the mode of action of RLs in plant defense. In order to validate the potential of RLs in plant protection against PPNs, their impact should be evaluated with crop plants and other important PPN species under non-axenic conditions.

## Material and Method

### Biosurfactant production

HAAs, mRLs and dRLs were produced in three different recombinant *Pseudomonas putida* KT2440 strains using the rhl gene set originating from *Pseudomonas aeruginosa. P. putida* KT2440 pSB01 [45] was used for the production of HAAs while *P. putida* KT2440 SK4 was used for the production of mRLs [46] and *P. putida* KT2440 pWJ02 was used for the production of predominantly dRLs [45]. The respective HAAs and RLs were a mixture of congeners with variation in the chain length of the β-OH fatty acids.

The strains were routinely grown on Luria Bertani (LB) agar plates. Cultivation was performed in Fernbach flasks filled with 500 ml LB medium supplemented with 20 g/L glucose and incubated in a Minitron Shaking Incubator (Infors AG, Bottmingen, Switzerland) at 30°C, 200 rpm, and 25 mm shaking diameter. LB medium contained 10 g/L tryptone, 5 g/L yeast extract and 5 g/L NaCl at pH 7.2. LB agar plates additionally contained 8 g/L agar. The final medium was sterilized by autoclaving for 20 min at 121°C before adding glucose. For cultivation of *P. putida* KT2440 pSB01 and *P. putida* KT2440 pJW02, kanamycin with a concentration of 50 μg/ml was added to maintain selection pressure. Overnight cultures were used to inoculate main cultures by adjusting the optical density at 600 nm to 0.1.

### Biosurfactant quantification

For quantification of hydroxyfatty acids, HAAs and RLs, components of the samples were separated using reversed-phase chromatography as described by Tiso *et al*. [47], based on the method established by Behrens *et al*. [48]. As HPLC system a DIONEX UltiMate 3000 (Thermo Fisher Scientific, Inc., Waltham, MA, USA) composed of an LPG-3400SD pump, a WPS-3000 (RS) autosampler, a TCC-3000 (RS) column oven, and a Corona charged aerosol detector (CAD) was used. The detector was supplied with a continuous nitrogen stream by a Parker Balston NitroVap-1LV nitrogen generator (Parker Hannifin GmbH, Kaarst, Germany). A NUCLEODUR C18 Gravity column from Macherey-Nagel GmbH and Co. KG (Düren, Germany) with a precolumn cartridge of 4 mm length, a particle size of 3 μm, and dimensions of 150×4.6 mm was utilized to separate the biosurfactants. The gradient program started with 70% (v/v) acetonitrile and 30% (v/v) ultrapure water containing 0.2% (v/v) formic acid. The acetonitrile percentage was constantly increased until it reached 80% (v/v) at 8 min and then kept constant for 1 min before it was further increased to 100% (v/v) within 1 min. After 11 min of total running time, the acetonitrile percentage was decreased again back to 70% (v/v) within 1.5 min to retrieve starting conditions. The total analysis time was 15 min. The column oven temperature was set to 40°C and 5 μl of the sample was injected. The flow rate was set to 1 ml/min. Samples were prepared by centrifugation at 13,000 × g for 1 min, whereupon 400 μl of the cell-free supernatant was mixed with 400 μl acetonitrile, and the resulting mixture was vortexed. Subsequently, samples were incubated overnight at 4°C to facilitate precipitation of proteins and other residuals that might clog the column. Next, biomass and precipitated material were removed by centrifugation at 13,000 × g for 1 min. Before HPLC analysis, the supernatant was filtered with a Phenex-RC (regenerated cellulose) syringe filter (diameter, 4 mm; pore size, 0.2 μm; Phenomenex, Torrance, CA, USA).

### Biosurfactant purification

Purification of HAAs and RLs was based on a method described previously [49]. Prior to purification, cells and proteins were removed from the culture broth by addition of acetonitrile as specified above. Isolation of the biosurfactants from the supernatant was then performed by solid phase extraction (SPE) using a C18 derivatized silica-based adsorbent (AA12SA5, YMC Europe GmbH, Dinslaken, Germany) packed into a Bioline glass column (Knauer GmbH, Berlin, Germany). For desorption, an AZURA analytical pump P 6.1L (Knauer GmbH, Berlin, Germany) was employed to carry out a gradient elution, starting with 100% double-distilled water and ending with 100% pure ethanol. The flow rate was set to 1 ml/min and a total of 21 fractions of 10 ml each were collected using a 2111 Multirac fraction collector (LKB, Bromma, Sweden). Fractions containing the desired RL congeners were combined and the solvent was evaporated.

For subsequent chromatographic separation of single RL congeners, the dried products were resolved in a 50/50 (v/v) acetonitrile/water mixture. The separation was carried out by the use of a preparative HPLC system consisting of an AZURA analytical pump P 6.1L, an AZURA autosampler 3950 (both Knauer GmbH, Berlin, Germany), a SEDEX 58 LT-ELSD detector (SEDERE, Olivet, France), and a Foxy R1 fraction collector (Teledyne ISCO Inc., Lincoln, NE, USA). As centerpiece of this setup, a NUCLEODUR C18 HTec column from Macherey-Nagel GmbH and Co. KG (Düren, Germany) with a precolumn cartridge of 16 mm length, a particle size of 5 μm, and dimensions of 250×21 mm was employed. The flow rate was set to 10 ml/min and a total amount of 200 mg of HAAs, 100 mg of mRLs or 10 mg of dRLs was injected. Elution was carried out starting with 70% (v/v) acetonitrile and 30% (v/v) 0.2% (v/v) formic acid. For the separation of HAAs and mRLs, acetonitrile was constantly increased until it reached 85% (v/v) at 45 min and then further increased to 100% (v/v) within 5 min. After 56 min of total running time, the acetonitrile percentage was decreased back to 70% (v/v) within 2 min to retrieve starting conditions. The total running time was 60 min. For the separation of dRLs, acetonitrile percentage was kept constant for 55 min and then increased to 100% (v/v) within 3 min. After 61 min of total running time, the acetonitrile percentage was decreased back to 70% (v/v) within 4 min to retrieve starting conditions. The total running time was 70 min. The desired RL congeners were fractionated according to peak elution, and subsequently evaporated to obtain pure, solvent-free biosurfactants. The dried products were solved in ultrapure water and the pH was adjusted to 7.0 by the use of 0.1 M NaOH and 0.1 M HCl. Finally, the preparations were filtered with a ROTILABO RC syringe filter (diameter 25 mm; pore size 0.2 μm; Carl Roth GmbH & Co. Kg, Karlsruhe, Germany) to guarantee sterility.

The final mRL mixture contained 10% C10-C8 mRLs, 62% C10-C10 mRL, 17% C10-C12 mRLs, 12% C10-C12:1 mRLs. The final dRL was comprised of 5% C10-C8 dRLs, 61% C10-C10 dRLs, 19% C10-C12 dRLs, 14% C10-C12:1 dRLs (Supplementary Figure 1). In all our experiments ppm corresponds to mg/l.

### Nematode mortality assay

A transparent flat bottom 96-well cell culture plate was prepared by filling a minimum of four wells per variant and time point with 100 μl dRLs (188.75, 377.5, 755, and 1510 ppm) or double distilled water as control further containing about 50 J2s prepared following Matera et al. [50]. Incubation was performed at room temperature and mortality was evaluated every 24 hours for four days using 10 μl of 5M sodium hydroxide (NaOH) per well with four technical replicates per time point and treatment. NaOH provokes movement of live nematodes and thus enables to reliably distinguish between alive and dead nematodes [50].

### Phenotyping of host-pathogen interaction

Seeds of *A. thaliana* wild type Col-0 were sterilized using 70% ethanol for 30 seconds followed by 1.3% sodium hypochlorite for three minutes. Sterile seeds were immediately washed three times in sterile double distilled water and dried onto sterile filter paper. For stratification, seeds were stored at 4°C until further use. Sowing was performed with five seeds per Petri dish filled with Knop agar [15] and stored at room temperature with 16/8 photoperiod in a tilted position to promote root surface growth for a facilitated transfer eight days post sowing (DPS). For the transfer, the plants were separated and two well- and similarly developed plants were transferred onto a new plate filled with about 16 ml RL-supplemented Knop agar or standard Knop agar as control. After two days of adaption, per plant 60-70 freshly hatched *H. schachtii* J2s [50] were inoculated in two drops along each primary root. Infection evaluation started 12 days post inoculation (DPI) by counting female and male nematodes under Stereo Microscope (Leica, Germany), followed by female size measurement 14 and 28 DPI using the LAS software (Leica Microsystems), and eventually female collection 35 DPI to invasively count the eggs and juveniles. The plant phenotype was ascertained 14 DPI by first separating shoot and root tissue at the hypocotyl and immediately weighing the fresh weight (FW) of the shoot with a fine balance. In order to remove the complete root system without breakage the agar was first dissolved and the agar-free root system was transferred into a transparent tray designed for subsequent root scanning (Epson Perfection V700 Photo root scanner) connected to the WinRHIZO Pro software (Regent Instruments, Québec, Canada) for automatic computation of the total root length.

### Reactive oxygen species analysis

*A. thaliana* seeds were sterilized and cultivated on standard Knop agar as described above, but with four seeds per plate. The Petri dishes were stored in a tilted position to ensure surface growth of the roots to facilitate damage-free root removal. About two weeks old leaves were harvested by cutting at the leaf base ensuring a similar size between all plant leaves and optimal fit into the well of a 96-well plate while the whole root system was cut at the hypocotyl. Plant tissue was incubated in sterile distilled water overnight in the dark the evening prior to the measurement to diminish the initial plant stress level. The following morning, a white flat bottom 96-well plate was filled with 80 μl of 100 μM L-012 sodium salt per well. Leaves were carefully transferred using a fine brush followed by another hour incubation in the dark. Prior to the measurement each well was further filled with 20 μl of horseradish peroxidase (HRP, 20 μg/ml) and 50 μl of either the test solution as threefold concentration or sterile double distilled water as negative control and 1 μM of the flg22 as positive control. The 96-well plate was instantly loaded into a microplate reader (Infinite ^®^ 200 PRO by Tecan Life Sciences Home, Germany) and luminescence was measured as relative light unit in a two-minute interval over a period of two hours.

### Gene expression analysis

*A. thaliana* wild type Col-0 seeds were sterilized and cultivated on standard Knop agar as described above, but with four seeds per plate. After eight days well-developed plants were transferred onto Knop agar supplemented with either 8.3 ppm mRLs or dRLs. The root system was sampled 1 hour and 48 hours after transplanting and transferred into a 1.5 ml Eppendorf tube for immediate freezing in liquid nitrogen and subsequent storage at −80°C until further use. Root total RNA was extracted with the RNeasy Plant Mini Kit (QIAGEN, Germany), genomic DNA contamination was removed with TURBO DNA-free™ Kit (Invitrogen, Germany) und reverse transcription was performed with the High-Capacity cDNA Reverse Transcription Kit (Thermofisher, Germany) and 100 ng RNA per reaction (Thermofisher) according to the respective manufacturer’s instructions. RT-PCR implied 20 μl per reaction with 10 μl 2X SYBR™ Green PCR Master Mix (Thermofisher), 0.5 μl of forward and reverse primer each (10 μM) and 1 μl cDNA (diluted 1:5) filled up with molecular grade water (supplementary table 1). Expression values were normalized to the expression of UBIQUITIN-SPECIFIC PROTEASE 22 (UBP22). One root system represented one biological replicate and was measured as three technical replicates.

### Statistical analysis

Statistical analysis and graphs were compiled with SigmaPlot (Version 12.5, Systat Software GmbH; Germany). For each experiment, the data of three independent biological repetitions were analysed altogether. Normality was tested based on Shapiro-Wilk with a P-value to reject of 0.05. Significance of multi comparison analyses was calculated based on ANOVA on ranks and a suitable post-hoc test. The Student Newman-Keuls (SNK) method was the post-hoc analysis of choice for experiments with an equal number of technical replicates while Dunn’s method was used for an unequal number among variants. Fold changes in root gene expression were normalized to the expression of the control. Significance in root gene expression was analysed based on an unpaired t-test in comparison to the control.

## Notes

### Competing Interest Statement

The authors have declared no competing interest.

