## Supplementary for "The biological activity of bacterial rhamnolipids is linked to their molecular structure"

**a**

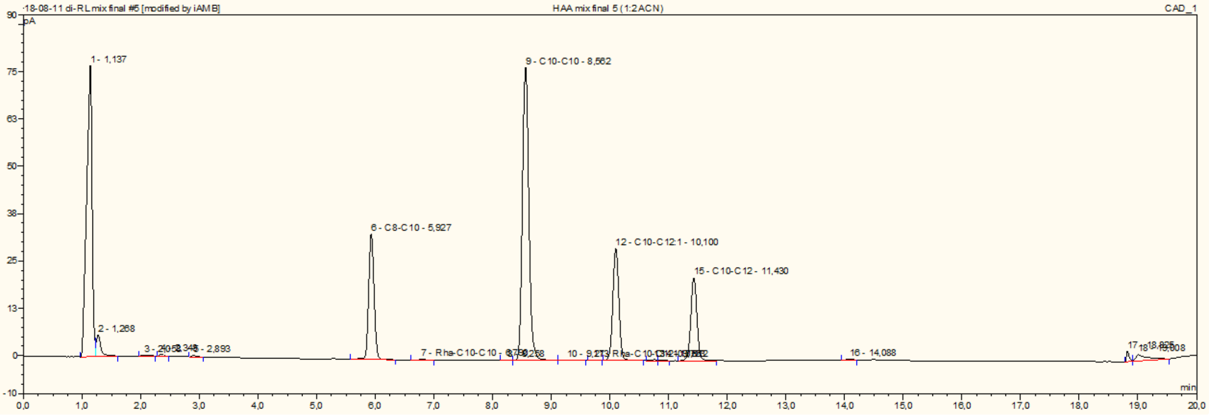


| | HAA | | | | mRL | | | | dRL | | | | HAA | mRL | dRL | surf. |
|---|---|---|---|---|---|---|---|---|---|---|---|---|---|---|---|---|
| | C8-C10 | C10-C10 | C10-C12:1 | C10-C12 | C8-C10 | C10-C10 | C10-C12:1 | C10-C12 | C8-C10 | C10-C10 | C10-C12:1 | C10-C12 | total | total | total | total |
| c [g/l] | 0.27 | 0.76 | 0.27 | 0.02 | - | - | - | - | - | - | - | - | 1.51 | - | - | 1.52 |

**b**

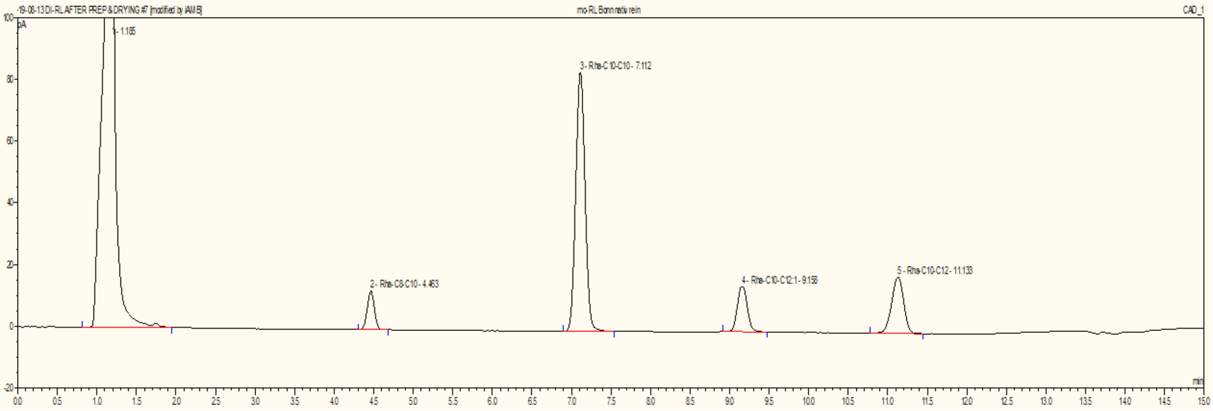


| | HAA | | | | mRL | | | | dRL | | | | HAA | mRL | dRL | surf. |
|---|---|---|---|---|---|---|---|---|---|---|---|---|---|---|---|---|
| | C8-C10 | C10-C10 | C10-C12:1 | C10-C12 | C8-C10 | C10-C10 | C10-C12:1 | C10-C12 | C8-C10 | C10-C10 | C10-C12:1 | C10-C12 | total | total | total | total |
| c [g/l] | - | - | - | - | 0.14 | 0.91 | 0.17 | 0.25 | - | - | - | - | - | 1.47 | - | 1.47 |

**c**

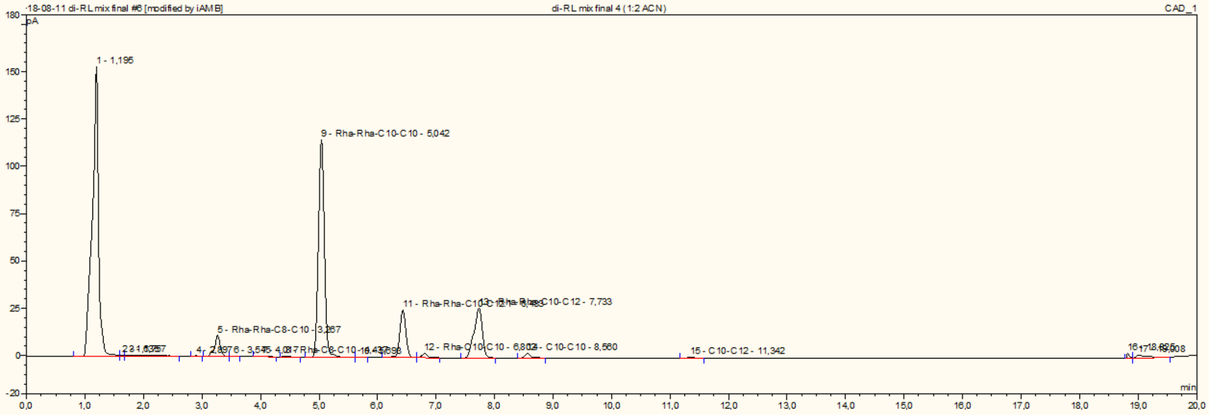


| | HAA | | | | mRL | | | | dRL | | | | HAA | mRL | dRL | surf. |
|---|---|---|---|---|---|---|---|---|---|---|---|---|---|---|---|---|
| | C8-C10 | C10-C10 | C10-C12:1 | C10-C12 | C8-C10 | C10-C10 | C10-C12:1 | C10-C12 | C8-C10 | C10-C10 | C10-C12:1 | C10-C12 | total | total | total | total |
| c [g/l] | - | 0.02 | - | - | - | 0.02 | - | - | 0.08 | 0.90 | 0.20 | 0.28 | 0.03 | 0.02 | 1.47 | 1.51 |

**d**

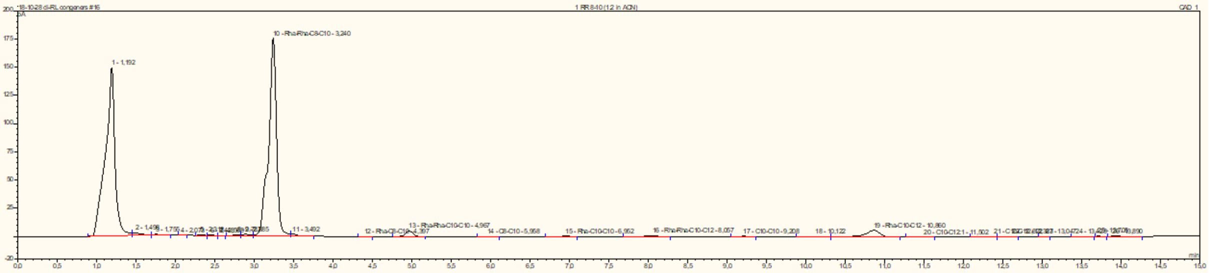

| | HAA | | | | mRL | | | | dRL | | | | HAA | mRL | dRL | surf. |
|---|---|---|---|---|---|---|---|---|---|---|---|---|---|---|---|---|
| | C8-C10 | C10-C10 | C10-C12:1 | C10-C12 | C8-C10 | C10-C10 | C10-C12:1 | C10-C12 | C8-C10 | C10-C10 | C10-C12:1 | C10-C12 | total | total | total | total |
| c [g/l] | - | - | - | - | - | - | - | - | 1.37 | - | - | - | - | - | 1.37 | 1.38 |

**e**

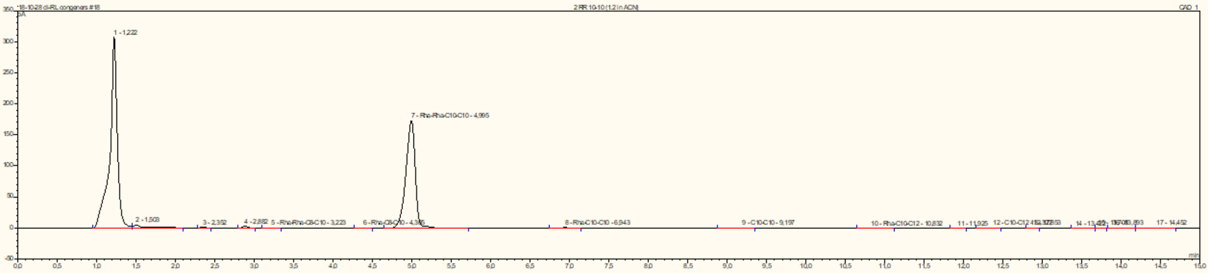

| | HAA | | | | mRL | | | | dRL | | | | HAA | mRL | dRL | surf. |
|---|---|---|---|---|---|---|---|---|---|---|---|---|---|---|---|---|
| | C8-C10 | C10-C10 | C10-C12:1 | C10-C12 | C8-C10 | C10-C10 | C10-C12:1 | C10-C12 | C8-C10 | C10-C10 | C10-C12:1 | C10-C12 | total | total | total | total |
| c [g/l] | - | - | - | - | - | - | - | - | - | 1.57 | - | - | - | - | 1.57 | 1.58 |

**f**

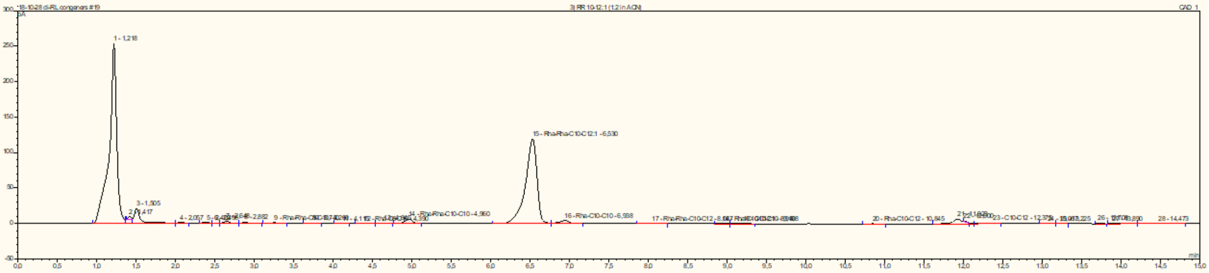

| | HAA | | | | mRL | | | | dRL | | | | HAA | mRL | dRL | surf. |
|---|---|---|---|---|---|---|---|---|---|---|---|---|---|---|---|---|
| | C8-C10 | C10-C10 | C10-C12:1 | C10-C12 | C8-C10 | C10-C10 | C10-C12:1 | C10-C12 | C8-C10 | C10-C10 | C10-C12:1 | C10-C12 | total | total | total | total |
| c [g/l] | - | - | - | - | - | - | - | - | - | - | 1.41 | - | - | - | 1.41 | 1.42 |

**g**

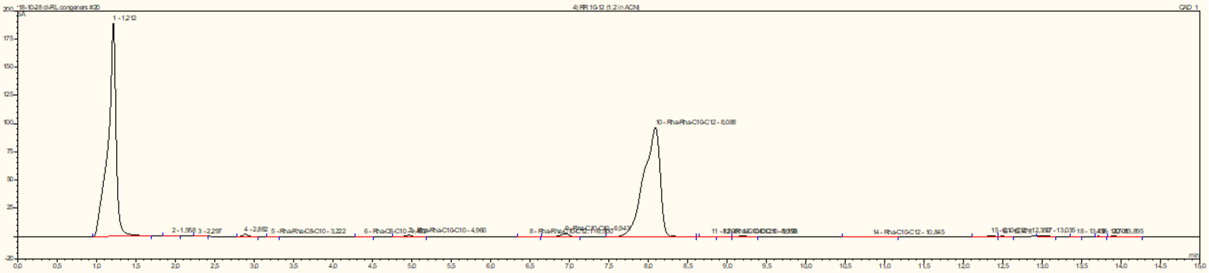

| | HAA | | | | mRL | | | | dRL | | | | HAA | mRL | dRL | surf. |
|---|---|---|---|---|---|---|---|---|---|---|---|---|---|---|---|---|
| | C8-C10 | C10-C10 | C10-C12:1 | C10-C12 | C8-C10 | C10-C10 | C10-C12:1 | C10-C12 | C8-C10 | C10-C10 | C10-C12:1 | C10-C12 | total | total | total | total |
| c [g/l] | - | - | - | - | - | - | - | - | - | - | - | 1.55 | - | - | 1.55 | 1.55 |

*Figure 1 Rhamnolipid composition. a) HAA mixture analogous to native production of P. aeruginosa PAO1. b) Mono-rhamnolipid (mRL) mixture analogous to native production of P. aeruginosa PAO1. c) di-rhamnolipid (dRL) mixture analogous to native production of P. aeruginosa PAO1. d) C10-C8 dRLs. e) C10-C10 dRLs. f) C10-C12:1 dRLs. g) C10-C12 dRLs.*

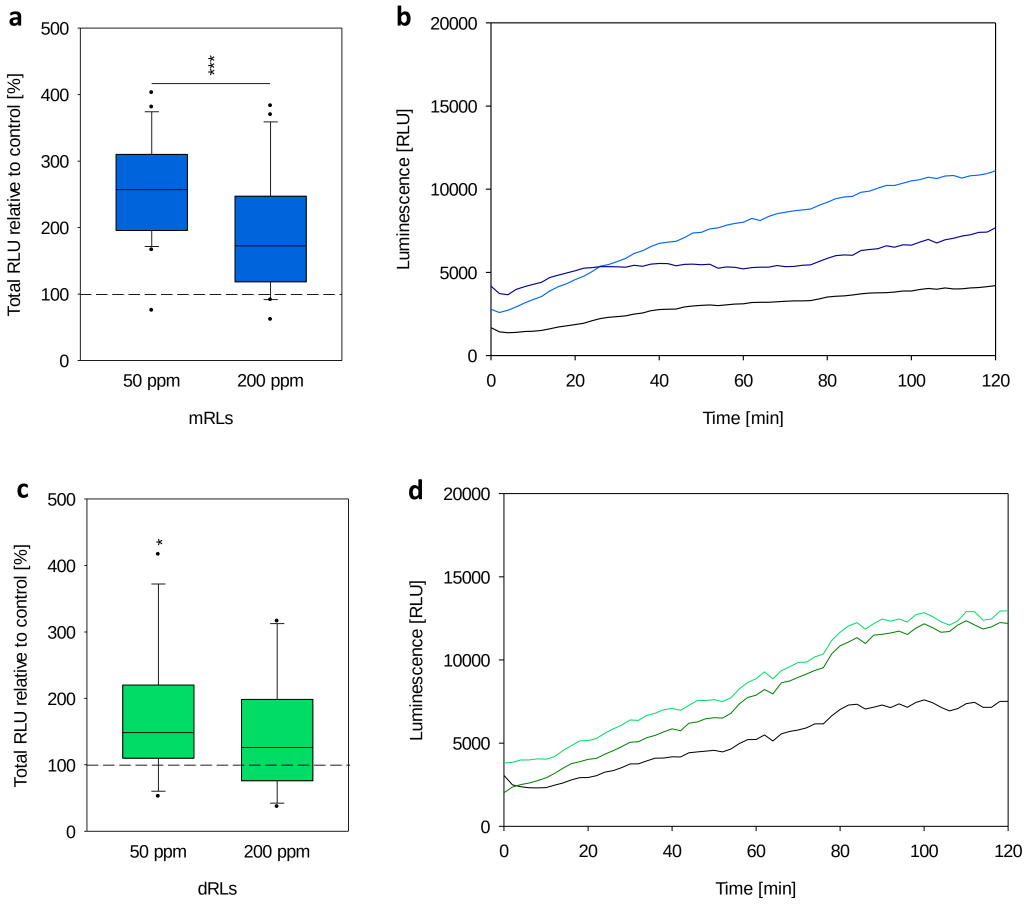


*Figure 2. Impact of different RLs on plant $H_2O_2$ production. Arabidopsis thaliana leaves were treated with mRLs and dRLs at different concentrations and subsequent synthesis of $H_2O_2$ was monitored over a period of two hours. a) Total $H_2O_2$ produced within 2 hours incubation in mRLs, median with 25%- and 75%- quartile, significance based on 2-tailed t-test compared to water control (dashed line) with * $p<0.05$, ** $p<0.01$, *** $p<0.001$. b) $H_2O_2$ production over time during mRL incubation, black: response to $ddH_2O$, from light to dark blue illustrates an increase in concentration (50, 200 ppm) of mRLs, mean values. c) Total $H_2O_2$ produced within 2 hours incubation in dRLs, median with 25%- and 75%- quartile, significance based on 2-tailed t-test compared to water control (dashed line) with * $p<0.05$, ** $p<0.01$, *** $p<0.001$. d) $H_2O_2$ production over time during dRL incubation, black: response to $ddH_2O$, from light to dark green illustrates an increase in concentration (50, 200 ppm) of dRLs, mean values. a)-d) Data of three independent biological replicates with total $n\geq11$.*

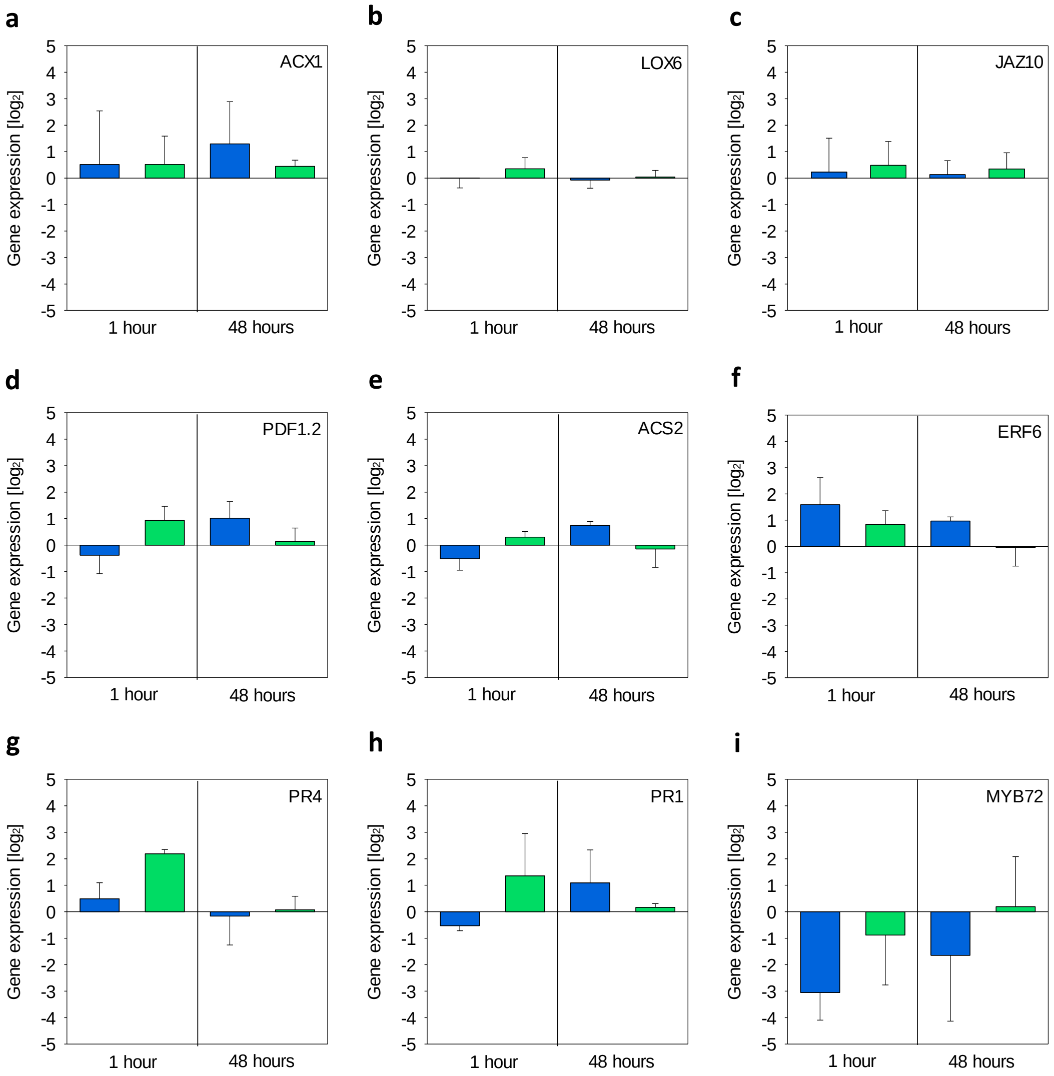


*Figure 3. Fold changes ($log_2$) in root gene expression of A. thaliana wild type after exposure to 8.3 ppm mono-rhamnolipids (mRLs, blue) or di-rhamnolipids (dRLs, green) for 1 or 48 hours in comparison to control plants on standard growth medium. a-c) Jasmonic acid (JA) defense gene markers. d) JA/ethylene (ET) defense gene marker. e-g) ET defense gene markers. h) Salicylic acid (SA) defense gene marker. i) Transcription factor for induced systemic resistance (ISR). Mean±SD of three independent biological replicates with 2≤n≤3, no significant differences.*

*Table 1 Selected Primer*

| | Sequence | Primer efficiency |
|---|---|---|
| ACX1 | 5′-CAT TTC GAT TTG TTG GAG AGT GGC TAA AA-3′<br>5′-AAC TCT TCA GAG CCT AGC GGT ACG AAG TT-3′ | 2.02 |
| ERF6 | 5'-GAA AAC CGC CGT TGA AGA TC-3'<br>5'-CGG TTG CGA ATT GAA TCC A-3' | 2.01 |
| PR1 | 5'-AAC TAC GCT GCG AAC ACG TG-3'<br>5'-TCA CTT TGG CAC ATC CGA GTC-3' | 2.06 |
| PR4 | 5'-AAC AAT GCG GTC AAG GC-3'<br>5'-AAG CAC TCA CGG CTC TCA AAT CCC-3' | 2 |
| PDF1.2 | 5'-CGC ACC GGC AAT GGT GGA AG-3'<br>5'-CAC ACG ATT TAG CAC CAA AG-3' | 1.91 |
| LOX6 | 5'-CCT CAT GAG AGA ATT TAT GAC TAC G-3'<br>5'-TGC TCT GAA GGT ATC TCT CTT GAT T-3' | 1.86 |
| JAZ10 | 5'-TCG CAA GGA GAA AGT CAC TGC AAC-3'<br>5'-CGA TTT AGC AAC GAC GAA GAA GGC-3' | 1.97 |
| ACS2 | 5'-GGA TGG TTT AGG ATT TGC TTT G-3'<br>5'-GCA CTC TTG TTC TGG ATT ACC TG-3' | 1.84 |
| MYB72 | 5’-CAA GCT TGC TGG ATT GTT GA-3‘<br>5’-AAG AAG GAC GCG ATC TTT GA-3‘ | 2.3 |
